# On the optimal trimming of high-throughput mRNA sequence data

**DOI:** 10.1101/000422

**Authors:** Matthew D. MacManes

**Affiliations:** University of New Hampshire. Durham, NH 03824; Department of Molecular, Cellular & Biomedical Sciences Durham, NH 03824; Hubbard Center for Genome Studies Durham, NH 03824

## Abstract

The widespread and rapid adoption of high-throughput sequencing technologies has afforded researchers the opportunity to gain a deep understanding of genome level processes that underlie evolutionary change, and perhaps more importantly, the links between genotype and phenotype. In particular, researchers interested in functional biology and adaptation have used these technologies to sequence mRNA transcriptomes of specific tissues, which in turn are often compared to other tissues, or other individuals with different phenotypes. While these techniques are extremely powerful, careful attention to data quality is required. In particular, because high-throughput sequencing is more error-prone than traditional Sanger sequencing, quality trimming of sequence reads should be an important step in all data processing pipelines. While several software packages for quality trimming exist, no general guidelines for the specifics of trimming have been developed. Here, using empirically derived sequence data, I provide general recommendations regarding the optimal strength of trimming, specifically in mRNA-Seq studies. Although very aggressive quality trimming is common, this study suggests that a more gentle trimming, specifically of those nucleotides whose Phred score *<*2 or *<*5, is optimal for most studies across a wide variety of metrics.

## Introduction

The popularity of genome-enabled biology has increased dramatically over the last few years. While researchers involved in the study of model organisms have had the ability to leverage the power of genomics for nearly a decade, this power is only now available for the study of non-model organisms. For many, the primary goal of these newer works is to better understand the genomic underpinnings of adaptive (Linnen et al., 2013; Narum et al., 2013) or functional (Hsu et al., 2012; Muñoz-Mérida et al., 2013) traits. While extremely promising, the study of functional genomics in non-model organisms typically requires the generation of a reference transcriptome to which comparisons are made. Although compared to genome assembly transcriptome assembly is less challenging (Bradnam et al., 2013; Earl et al., 2011), significant computational hurdles still exist. Amongst the most difficult of challenges in transcriptome assembly involves the reconstruction of isoforms (Pyrkosz et al., 2013), simultaneous assembly of transcripts where read coverage (=expression) varies by orders of magnitude, and overcoming biases related to random hexamer (Hansen et al., 2010) and GC content (Dohm et al., 2008).

These processes are further complicated by the error-prone nature of high-throughput sequencing reads. With regards to Illumina sequencing, error is distributed non-randomly over the length of the read, with the rate of error increasing from 5’ to 3’ end (Liu et al., 2012). These errors are overwhelmingly substitution errors (Yang et al., 2013), with the global error rate being between 1% and 3%. Although *de Bruijn* graph assemblers do a remarkable job in distinguishing error from correct sequence, sequence error does results in assembly error (MacManes and Eisen, 2013). While this type of error is problematic for all studies, it may be particularly troublesome for SNP-based population genetic studies. In addition to the biological concerns, sequencing read error may results in problems of a more technical importance. Because most transcriptome assemblers use a *de Bruijn* graph representation of sequence connectedness, sequencing error can dramatically increase the size and complexity of the graph, and thus increase both RAM requirements and runtime.

In addition to sequence error correction, which has been shown to improve accuracy of the *de novo* assembly (MacManes and Eisen, 2013), low quality (=high probability of error) nucleotides are commonly removed from the sequencing reads prior to assembly, using one of several available tools (Trimmomatic (Lohse et al., 2012), Fastx Toolkit (http://hannonlab.cshl.edu/fastx_toolkit/index.html), or biopieces (http://www.biopieces.org/). These tools typically use either a sliding window approach, discarding nucleotides falling below a given (user selected) average quality threshold, or trimming of low-quality nucleotides at one or both ends of the sequencing read. Though the absolute number will surely be decreased in the trimmed dataset, aggressive quality trimming may remove a substantial portion of the total read dataset, which in transcriptome studies may disproportionately effect lower expression transcripts.

Although the process of nucleotide quality trimming is commonplace, particularly in the assembly-based HTS analysis pipelines (e.g. SNP development (Helyar et al., 2012; Milano et al., 2011), functional studies (Ansell et al., 2013; Bhardwaj et al., 2013), and more general studies of transcriptome characterization (Liu et al., 2013; MacManes and Lacey, 2012)), its optimal implementation has not been well defined. Though the rigor with which trimming is performed may be guided by the design of the experiment, a deeper understanding of the effects of trimming is desirable. As transcriptome-based studies of functional genomics continue to become more popular, understanding how quality trimming of mRNA-seq reads used in these types of experiments is urgently needed. Researchers currently working in these field appear to favor aggressive trimming (e.g. (Looso et al., 2013; Riesgo et al., 2012)), but this may not be optimal. Indeed, one can easily image aggressive trimming resulting in the removal of a large amount of high quality data (even nucleotides removed with the commonly used Phred=20 threshold are accurate 99% of the time), just as lackadaisical trimming (or no trimming) may result in nucleotide errors being incorporated into the assembled transcriptome.

Here, to provide recommendations regarding the efficient trimming of high-throughput sequence reads, specifically for mRNASeq reads from the Illumina platform. To do this, I used publicly available datasets containing Illumina reads derived from *Mus musculus*. Subsets of these data (10 million, 20 million, 50 million, 75 million, 100 million reads) were randomly chosen, trimmed to various levels of stringency, assembled then analyzed for assembly error and content. In addition to this, I develop a set of metrics that may be generally useful in evaluating the quality of transcriptome assemblies. These results aim to guide researchers through this critical aspect of the analysis of high-throughput sequence data. While the results of this paper may not be applicable to all studies, that so many researchers are interested in the genomics of adaptation and phenotypic diversity, particularly in non-model organisms suggests its widespread utility.

## Materials and Methods

Because I was interested in understanding the effects of sequence read quality trimming on the quality of vertebrate transcriptome assembly, I elected to analyze a publicly available (SRR797058) paired-end Illumina read dataset. This dataset is fully described in a previous publication (Han et al., 2013), and contains 232 million paired-end 100nt Illumina reads. To investigate how sequencing depth influences the choice of trimming level, reads data were randomly subsetted into 10 million, 20 million, 50 million, 75 million, 100 million read datasets. To test the robustness of my findings, I evaluated a second dataset (SRR385624, (Macfarlan et al. (2012)) as well as a technical replicate of the primary dataset, both at the 10M read dataset size.

Read datasets were trimmed at varying quality thresholds using the software package Trimmomatic version 0.30 (Lohse et al., 2012), which was selected as it appears to be amongst the most popular of read trimming tools. Specifically, sequences were trimmed at both 5’ and 3’ ends using Phred =0 (adapter trimming only), *≤* 2, *≤* 5, *≤* 10, and *≤* 20. Other parameters (MINLEN=25, ILLUMINACLIP=barcodes.fa:2:40:15, SLIDINGWINDOW size=4) were held constant. Transcriptome assemblies were generated for each dataset using the default settings (except group_pairs_distance flag set to 999) of the program Trinity r2013-02-25 (Grabherr et al., 2011; Haas et al., 2013). Assemblies were evaluated using a variety of different metrics, many of them comparing assemblies to the complete collection of *Mus* cDNA’s, available at http://useast.ensembl.org/info/data/ftp/index.html.

Quality trimming may have substantial effect on assembly quality, and as such, I sought to identify high quality transcriptome assemblies. Assemblies with few nucleotide errors relative to a known reference may indicate high quality. The program Blat v34 (Kent, 2002) was used to identify and count nucleotide mismatches between reconstructed transcripts and their corresponding reference. To eliminate spurious short matches between query and template inflating estimates of error, only unique transcripts that covered more than 90% of their reference sequence were used. Next, because kmers represent the fundamental unit of assembly, kmers (k=25) were counted for each dataset using the program Jellyfish v1.1.11 (Marçais and Kingsford, 2011). Another potential assessment of assembly quality may be related to the number of paired-end sequencing reads that concordantly map to the assembly. As the number of reads concordantly mapping increased, so does assembly quality. To characterize this, I mapped the full dataset (not subsampled) of adapter trimmed sequencing reads to each assembly using Bowtie2 v2.1.0 (Trapnell et al., 2010) using default settings, except for maximum insert size (-X 999) and number of multiple mappings (-k 30).

Aside from these metrics, measures of assembly content were also assayed. Here, open reading frames (ORFs) were identified using the default settings of the program TransDecoder r20131110 (http://transdecoder.sourceforge.net/), and were subsequently translated into amino acid sequences, both using default settings. The larger the number of complete open reading frames (containing both start and stop codons) the better the assembly. Next, unique transcripts were identified using the blastP program within the Blast+ package version 2.2.28 (Camacho et al., 2009). Blastp hits were retained only if the sequence similarity was *>*80% over at least 100 amino acids, and e-value <10^−10^. As the number of transcripts matching a given reference increases, so may assembly quality. Lastly, because the effects of trimming may vary with expression, I estimated expression (e.g. FPKM) for each assembled contig using default settings of the the program eXpress v1.5.0 (Roberts and Pachter, 2013) and the BAM file produced by Bowtie2 as described above. Code for performing the subsetting, trimming, assembly, peptide and ORF prediction and blast analyses can be found in the following Github folder https://github.com/macmanes/trimming_paper/tree/recreate_ms_analyses/scripts.

## Results

Quality trimming of sequence reads had a relatively large effect on the total number of errors contained in the final assembly (Figure 1), which was reduced by between 9 and 26% when comparing the assemblies of untrimmed versus Phred=20 trimmed sequence reads. Most of the improvement in accuracy is gained when trimming at the level of Phred=5 or greater, with modest improvements potentially garnered with more aggressive trimming at certain coverage levels (Table 1).

**Figure 1.**
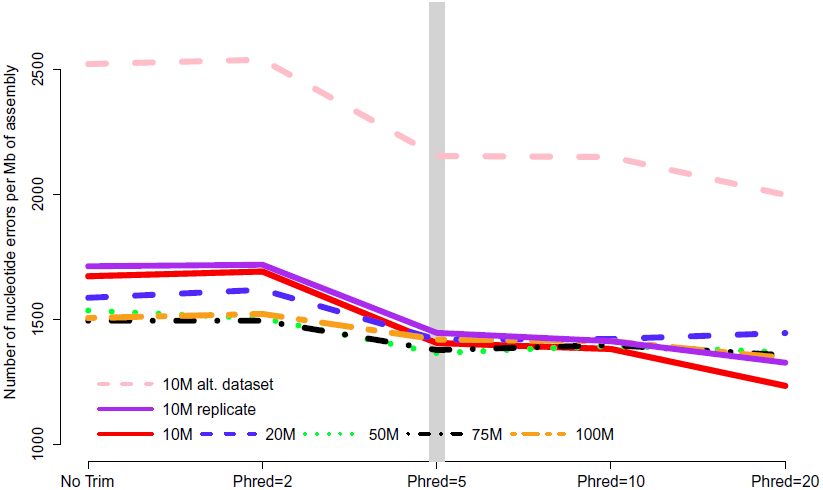
The number of nucleotide errors contained in the final transcriptome assembly, normalized to assembly size, is related to the strength of quality trimming. This patterns is largely unchanged with varying depth of sequencing coverage (10 million to 100 million sequencing reads). Trimming at Phred = 5 may be optimal, given the potential untoward effects of more stringent quality trimming. 10M, 20M, 50M, 75M, 100M refer to the subsamples size. 10M replicate is the technical replicate, 10M alt. dataset is the secondary dataset. Note that to enhance clarity, the Y-axis does not start at zero.

**Table 1.** The Phred trimming levels that resulted in optimal assemblies across the 4 metrics tested in the different size datasets. Error= the number of nucleotide errors in the assembly. Map= the number of concordantly mapped reads. ORF= the number of ORFs identified. BLAST= the number of unique BLAST hits. rep. is the technical replicate, 10M alt. is the secondary dataset.

| DATASET SIZE | ERROR | MAP | ORF | BLAST |
| --- | --- | --- | --- | --- |
| 10M | 20 | 0 | 2 | 2 |
| 10M REP. | 20 | 2 | 2 | 2 |
| 10M ALT | 20 | 2 | 0 | 0 |
| 20M | 5 | 5 | 2 | 2 |
| 50M | 5 | 10 | 5 | 2 |
| 75M | 20 | 10 | 5 | 0 |
| 100M | 20 | 0 | 2 | 2 |

In *de Bruijn* graph-based assemblers, the kmer is the fundamental unit of assembly. Even in transcriptome datasets, unique kmers are likely to be formed as a results of sequencing error, and therefore may be removed during the trimming process. Figure 2A shows the pattern of unique kmer loss across the various trimming levels and read datasets. What is apparent, is that trimming at Phred=5 removes a large fraction of unique kmers, with either less- or more-aggressive trimming resulting in smaller effects. In contrast to the removal of unique kmers, those kmers whose frequency is >1 are more likely to be real, and therefore should be retained. Figure 2B shows that while Phred=5 removes unique kmers, it may also reduce the number of non-unique kmers, which may hamper the assembly process.

**Figure 2A.**
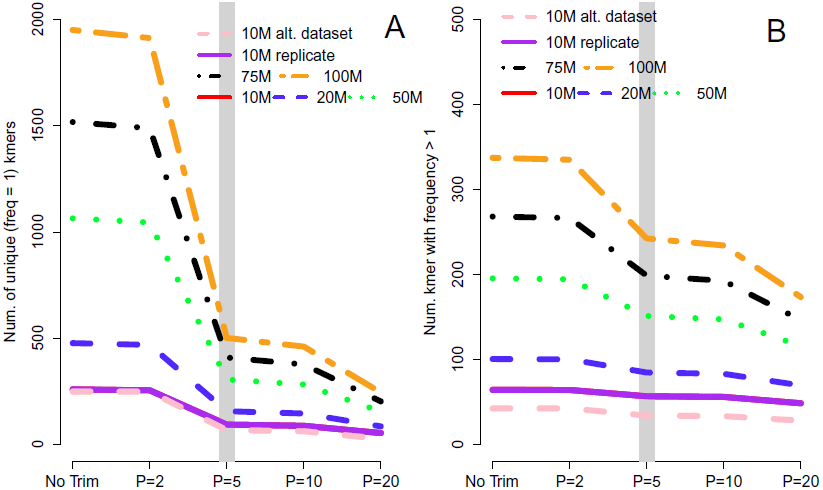
The number of unique kmers removed with various trimming levels across all datasets. Trimming at Phred=5 results in a substantial loss of likely erroneous kmers, while the effect of more and less aggressive trimming is more diminished. 2B depicts the relationship between trimming and non-unique kmers, whose pattern is similar to that of unique kmers.

In addition to looking at nucleotide error and kmer distributions, assembly quality may be measured by the the proportion of sequencing reads that map concordantly to a given transcriptome assembly (Hunt et al., 2013). As such, the analysis of assembly quality includes study of the mapping rates. Here, I found small but important effects of trimming. Specifically, assembling with aggressively quality trimmed reads decreased the proportion of reads that map concordantly. For instance, the percent of reads successfully mapped to the assembly of 10 million Q20 trimmed reads was decreased by 0.6% or approximately 1.4 million reads (compared to mapping of untrimmed reads) while the effects on the assembly of 100 million Q20 trimmed reads was more blunted, with only 381,000 fewer reads mapping. Though the differences in mapping rates are exceptionally small, when working with extremely large datasets, the absolute difference in reads utilization may be substantial.

Analysis of assembly content painted a similar picture, with trimming having a relatively small, though tangible effect. The number of BLAST+ matches decreased with stringent trimming (Figure 3), with trimming at Phred=20 associated with particularly poor performance. The maximum number of BLAST hits for each dataset were 10M=27452 hits, 20M=29563 hits, 50M=31848 hits, 75M=32786 hits, and 100M=33338 hits.

**Figure 3.**
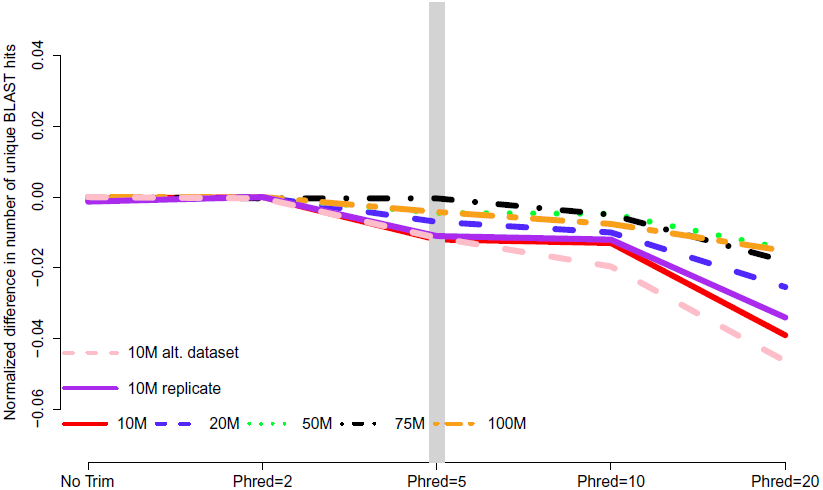
The number of unique Blast matches contained in the final transcriptome assembly is related to the strength of quality trimming, with more aggressive trimming resulting in worse performance. Data are normalized to the number of BLAST hits obtained in the most favorable trimming level for each dataset. Negative numbers indicate the detrimental affect of trimming. 10M, 20M, 50M, 75M, 100M refer to the subsamples size. 10M replicate is the technical replicate, 10M alt. dataset is the secondary dataset.

When counting complete open reading frames recovered in the different assemblies, all datasets were all worsened by aggressive trimming, as evidenced by negative values in Figure 4. Trimming at Phred=20 was the most poorly performing level at all read depths. The maximum number of complete open reading frames for each dataset were 10M=11429 ORFs, 20M=19463 ORFs, 50M=35632 ORFs, 75M=42205 ORFs, 100M=48434 ORFs.

Of note, all assembly files will be deposited in Dryad upon acceptance for publication. Until then, they can be accessed via https://www.dropbox.com/sh/oiem0v5jgr5c5ir/TYQdGcpYwP.

**Figure 4.**
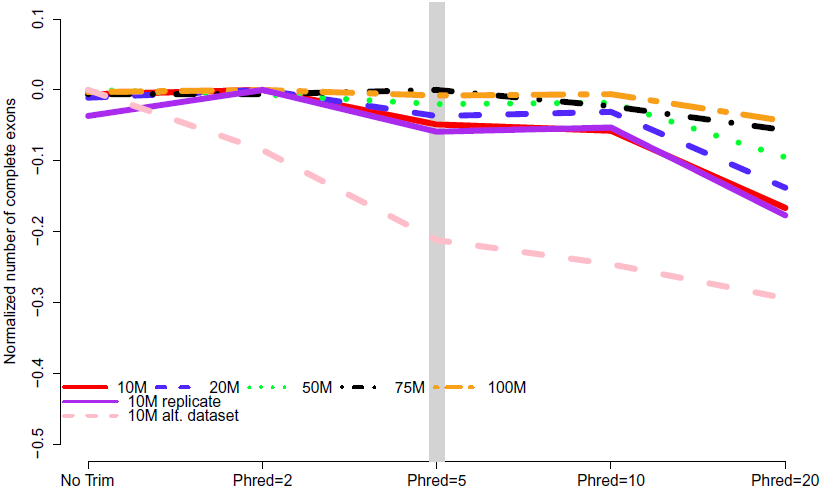
The number of complete exons contained in the final transcriptome assembly is related to the strength of quality trimming for any of the studied sequencing depths, Trimming at Phred=20 was always associated with poor performance. Data are normalized to the number of complete exons obtained in the most favorable trimming level for each dataset. Negative numbers indicate the detrimental affect of trimming. 10M, 20M, 50M, 75M, 100M refer to the subsamples size. 10M replicate is the technical replicate, 10M alt. dataset is the secondary dataset.

## Discussion

Although the process of nucleotide quality trimming is commonplace in HTS analysis pipelines, particularly those involving assembly, its optimal implementation has not been well defined. Though the rigor with which trimming is performed seems to vary, there is a bias towards stringent trimming (Ansell et al., 2013; Barrett and Davis, 2012; Straub et al., 2013; Tao et al., 2013). This study provides strong evidence that stringent quality trimming of nucleotides whose quality scores are *≤* 20 results in a poorer transcriptome assembly across the majority metrics. Instead, researchers interested in assembling transcriptomes *de novo* should elect for a much more gentle quality trimming, or no trimming at all. Table 1 summarizes my finding across all experiments, where the numbers represent the trimming level that resulted in the most favorable result. What is apparent, is that for typically-sized datasets, trimming at Phred=2 or Phred=5 optimizes assembly quality. The exception to this rule appears to be in studies where the identification of SNP markers from high (or very low) coverage datasets is the primary goal.

The results of this study were surprising. In fact, much of my own work assembling transcriptomes included a vigorous trimming step. That trimming had generally small effects, and even negative effects when trimming at Phred=20 was unexpected. To understand if trimming changes the distribution of quality scores along the read, we generated plots with the program SolexaQA (Cox et al., 2010). Indeed, the program modifies the distribution of Phred scores in the predicted fashion yet downstream effects are minimal. This should be interpreted as speaking to the performance of the the bubble popping algorithms included in Trinity and other *de Bruijn* graph assemblers.

The majority of the results presented here stem from the analysis of a single Illumina dataset and specific properties of that dataset may have biased the results. Though the dataset was selected for its ‘typical’ Illumina error profile, other datasets may produce different results. To evaluate this possibility, a second dataset was evaluated at the 10M subsampling level. Interestingly, although the assemblies based on this dataset contained more error (e.g. Figure 1), aggressive trimming did not improve quality for any of the assessed metrics, though like other datasets, the absolute number of errors were reduced.

In addition to the specific dataset, the subsampling procedure may have resulted in undetected biases. To address these concerns, a technical replicate of the original dataset was produced at the 10M subsampling level. This level was selected as a smaller sample of the total dataset is more likely to contain an unrepresentative sample than larger samples. The results, depicted in all figures as the solid purple line, are concordant. Therefore, I belive that sampling bias is unlikely to drive the patterns reported on here.

What is missing in trimmed datasets? — The question of differences in recovery of specific contigs is a difficult question to answer. Indeed, these relationships are complex, and could involve a stochastic process, or be related to differences in expression (low expression transcripts lost in trimmed datasets) or length (longer contigs lost in trimmed datasets). To investigate this, I attempted to understand how contigs recovered in the 10 million read untrimmed dataset, but not in the Phred=20 trimmed dataset were different. Using the information on FPKM and length generated by the program eXpress, it was clear that the transcripts unique to the untrimmed dataset were more lowly expressed (mean FPKM=3.2) when compared to the entire untrimmed dataset (mean FPKM=11.1; W = 18591566, p-value = 7.184e-13, non-parametric Wilcoxon test).

I believe that the untoward effects of trimming are linked to a reduction in coverage. For the datasets tested here, trimming at Phred=20 resulted in the loss of nearly 25% of the dataset, regardless of the size of the initial dataset. This relationship does suggest, however, that the magnitude of the negative effects of trimming should be reduced in larger datasets, and in fact may be completely erased with ultra-deep sequencing. Indeed, when looking at the differences in the magnitude of negative effects in the datasets presented here, it is apparent that trimming at Phred=20 is ‘less bad’ in the 100M read dataset than it is in the 10M read datasets. For instance, Figure 2B demonstrates that one of the untoward effects of trimming, the reduction of non-unique kmers, is reduced as the depth of sequencing is increased. Figures 3 and 4 demonstrate a similar pattern, where the negative effects of aggressive trimming of higher coverage datasets are blunted relative to lower coverage datasets.

Turning my attention to length, when comparing uniquely recovered transcripts to the entire untrimmed dataset of 10 million reads, it appears to be the shorter contigs (mean length 857nt versus 954nt; W = 26790212, p-value <2.2e-16) that are differentially recovered in the untrimmed dataset relative to the Phred=20 trimmed dataset.

Effects of coverage on transcriptome assembly — Though the experiment was not designed to evaluate the effects of sequencing depth on assembly, the data speak well to this issue. Contrary to other studies, suggesting that 30 million paired end reads were sufficient to cover eukaryote transcriptomes (Francis et al., 2013), the results of the current study suggest that assembly content was more complete as sequencing depth increased; a pattern that holds at all trimming levels. Though the suggested 30 million read depth was not included in this study, all metrics, including the number of assembly errors, as well as the number of exons, and BLAST hits were improved as read depth increased. While generating more sequence data is expensive, given the assembled transcriptome reference often forms the core of future studies, this investment may be warranted.

Should quality trimming be replaced by unique kmer filtering? — For transcriptome studies that revolve around assembly, quality control of sequence data has been thought to be a crucial step. Though the removal of erroneous nucleotides is the goal, how best to accomplish this is less clear. As described above, quality trimming has been a common method, but in its commonplace usage, may be detrimental to assembly. What if, instead of relying on quality scores, we instead rely on the distribution of kmers to guide our quality control endeavors? In transcriptomes of typical complexity, sequenced to even moderate coverage, it is reasonable to expect that all but the most exceptionally rare mRNA molecules are sequenced at a depth >1. Following this, all kmer whose frequency is <2 are putative errors, and should be removed before assembly, though this process may result in the loss of kmers from extremely low abundance transcripts or isoforms. This idea and its implementation are fodder for future research.

In summary, the process of nucleotide quality trimming is commonplace in many HTS analysis pipelines, but its optimal implementation has not been well defined. A very aggressive strategy, where sequence reads are trimmed when Phred scores fall below 20 is common. My analyses suggest that for studies whose primary goal is transcript discovery, that a more gentle trimming strategy (*e.g.* Phred=2 or Phred=5) that removes only the lowest quality bases is optimal. In particular, it appears as if the shorter and more lowly expressed transcripts are particularly vulnerable to loss in studies involving more harsh trimming. The one potential exception to this general recommendation may be in studies of population genomics, where deep sequencing is leveraged to identify SNPs. Here, a more stringent trimming strategy may be warranted.

## Acknowledgments

This paper was greatly improved by suggestions of C. Titus Brown and Christian Cole. In addition, the paper was first released as a bioRxiv preprint, and I received several comments based on that work both on that website as well as via Twitter. Let it be said here, that early use of a preprint archive, open access publication, and Twitter based discussion is a powerful way to rapidly disseminate (and get feedback on) work. I highly encourage its use!

